# Color traits of soybean conventional and trangenic cultivars are affected by glyphosate concentrations using digital imaging

**DOI:** 10.1101/2023.04.11.536485

**Authors:** Francisco Cleilson Lopes Costa, Mateus Ribeiro Piza, Lais Nóbrega Rodrigues, Adriano Teodoro Bruzi, Welison Andrade Pereira

## Abstract

Soybean is the most cultivated oilseed crop in the world, with much of the merit obtained in recent years due to genetic improvement in which greater genetic progress can be obtained by improving physiological characteristics, which result in a greater impact on yield. Color spectra can be good indicators of the physiological quality of plants by quantifying the intensity of colors in the RGB spectra, being a non-destructive method that allows optimizing the collection and the number of data. We aimed to identify differences in spectral reflection between conventional, transgenic RR1 and RR2 soybean cultivars under the effect of glyphosate herbicide. The plants were cultivated in five-liter pots in a greenhouse in a randomized block design, following cultural treatments recommended for the soybean crop. Leaves of plants in vegetative stages V1 or V2 were collected and duly identified to compose a second experiment, being then submitted to incubation in plastic trays in which the treatments were organized in a completely randomized design in a 6×6 factorial scheme, with three replications. Glyphosate herbicide doses (0.0, 0.03%, 0.06%, 0.12% and 0.24% AE/ha) were added to the leaf petiole on a cotton pad in order to maintain constant contact with the respective dose. The trays were kept under ambient conditions for 14 days after incubation under 16h-light and 8h-dark of artificial light. Images were collected with a smartphone camera on the 13th day after leaves collection (DAC) in an appropriate studio to maintain adequate lighting, and on the 14th DAC SPAD index data were collected in three locations on the leaf, avoiding the midrib. The images were cropped and then segmented using manual thresholding in which the average values for the red, green and blue channels were extracted. The Excess Red Index (ExR) was calculated using the red and green channels data. The data obtained were analyzed and the significant effects of the model were analyzed by the Skott-Knott test for the cultivar factor and regression models were adjusted for the dose factor. Spearman’s correlation was used to verify the relationship between the studied variables. In view of our results, glyphosate affects the chlorophyll of resistant plants when subjected to continuous exposure and at high doses, leading to senescence and that the red channel information can be used to infer the level of interference in the photosynthetic activity of plants subjected to the herbicide.

## Introduction

Soybean is the most cultivated oilseed crop in the world, with Brazil, USA and China being the largest producers reaching about 80% of the world production. Productivity is expected to continue increasing due to genetic improvement. Greater genetic progress can be achieved by improving physiological traits, which result in a greater impact on yield. Color is a good indicator of the physiological quality of plants. The use of herbicides in soybean crops can directly affect the physiology of plants, even of transgenic cultivars. Genetic improvement can help reduce this harmful effect of the herbicide on the crop. Therefore, quantifying the the digital numbers of the channels of the RGB spectra can be useful to adderss this task.

One of the physiological traits inherent to yield is the herbicide tolerance of RR cultivars. In a recent study, the authors demonstrated that the use of the glyphosate herbicide directly interferes with the translocation of dry matter, even in RR cultivars, impacting root development (COSTA; PEÑA; PEREIRA, 2022). One of the ways to approach and better understand the problem is to perform an further evaluation through images. Thus, questions arise about the existence of other associations between herbicide application and impacts on productivity, radiation use efficiency, and whether there are possibilities related to genetic improvement for the use of radiation under the effect of the herbicide.

Great advances in productivity were achieved without knowledge of the molecular mechanisms of the plant. Molecular genetics supports the understanding of the function of genes and physiological processes related to the productive potential of the crop. To our knowledge, no study in the literature exists regarding interaction with GMO soy and radiation, in terms of how photosynthetic pigments can be affected under the effect of glyphosate.

Chlorophylls strongly absorb radiation in the visible spectrum, resulting in low reflectance. However, leaves have a relatively high absorption in the near infrared due to internal leaf scattering and no reflectance, and in the infrared (>1.3 μm) reflectance is relatively low due to the absorption of radiation by the water (GATES et al., 1965; KNIPLING, 1970).

Continuous stress can harzardous to the plant leading to senescence, a process that involves endogenous physiological changes and exogenous stressors (CHEN, 2014), helping the plant to respond against biotic and abiotic stresses by recycling and reusing nutrients from senescent leaves in newly developed or developing organs, increasing the survival rate of plants (DANI et al., 2016; KIM; WOO; NAM, 2016). In plant development, leaf senescence is the last process, being regulated by a large number of genes. In rice, 185 genes are related to chloroplast degradation, energy metabolism routes, nitrogen mobilization, hormones and transcription factors (LIM; KIM; GIL NAM, 2007; LI et al., 2020).

For behavioral studies, it is ideal to monitor changes in the same leaf, to assess the effects of treatment, senescence, and abiotic stress over time. Measuring chlorophyll content is a slow chemical process that involves destruction of plant tissue, in addition to using toxic chemical reagents. Despite being an important indicator of physiological and health status, the measurement of photosynthetic capacity can be performed indirectly through digital images, adopting highly correlated characteristics.

Photosynthetic pigments such as chlorophylls a and b, xanthophylls and carotenes are components of photosystems I and II, essential for photosynthesis. Chlorophylls absorb two types of wavelengths in the visible light spectrum, starting to absorb in blue light. The main photoreceptors, such as cryptochromes (MISHRA; KHURANA, 2017), phototropins (MAO et al., 2005), UVR8 (CHRISTIE et al., 2012), zeitlupes (CHRISTIE et al., 2015), and phytochromes (CARVALHO; CAMPOS; AZEVEDO, 2013) that compose systems responsible for sensing variation in the light and adapting the plant to diverse environmental conditions.

There are many alternatives for monitoring chlorophyll content in leaves, however direct measurement of pigment concentration seems to be the most reliable method. However, indirect and non-destructive methods are highlighted to accomplish this task. Non-destructive methods are those that indirectly evaluate the chlorophyll content in a precise, economical way, with high profitability and quality, allowing the evaluation of many plants in a single evaluation.

One of these alternatives for non-destructive evaluation is through the SPADmeter, an equipment developed to provide a relative intensity of the green of the leaf based on two light emitting diodes and a silicon photodiode receiver (MINOLTA, 1989). The SPADmeter provides a score that is indicative of the chlorophyll content of the leaf, with a scale ranging from -9.99 to 199.99. Despite the great advantage presented by the SPADmetro, its evaluation is done sheet by sheet, which greatly reduces the evaluation yield, when thinking about large experiments in the field. To know the exact value of leaf chlorophyll concentration through SPAD, it is necessary to derive a calibration curve with SPAD values and the relative chlorophyll content (LI; AUBREY; SUN, 2017). Chlorophyll reduction is a great alternative to evaluate leaf senescence (MATILE et al., 1996), a great indicator for states of abiotic stress. Digital images are presented as a very useful resource to evaluate agricultural experiments, as they increase the efficiency of the evaluation, allowing obtaining several quantitative characteristics of the segmented image, in addition to being a non-destructive methodology (HATEM; TAN, 2003). The use of digital RGB images associated with analysis software has been used in applications, for example, of leaf senescence in wheat (ADAMSEN et al., 1999), quantification of canopy cover in wheat (LUKINA; STONE; RAUN, 1999) and soybean (PURCELL, 2000), coverage and senescence in a corn field (MAKANZA et al., 2018), phenotyping of energetic sorghum with a robot-coupled RGB camera (YOUNG; KAYACAN; PESCHEL, 2019).

In soybean culture, several research works were carried out, e.g., to evaluate the relationship between nitrogen fixation through comparative analyzes between the SPAD index and the evaluation of digital images to determine the chlorophyll content, Vollmann et al., (2011) evaluated nodulating and non-nodulating soybean lines and performed the phenotyping of the nodulation vs. non-nodulation that were associated with foliar, agronomic and seed characteristics. In addition, image data related to the green color value of the RGB space and chlorophyll (SPADmeter) were negatively correlated (R = -0.937) at the R3 developmental stage, when pod growth begins (VOLLMANN et al., 2011). Gwata (2004) carried out an experiment to indirectly determine leaf chlorophyll content in soybeans through a color score and reported that there is a correlation of 0.88 (df = 19, P < 0.01) between leaf chlorophyll content and the color score. Close correlation between parameters related to stress by nitrogen indicator by chlorophyll meter and information obtained from multispectral images has already been proven (REUM; ZHANG, 2007). Therefore, the aim of this study was to identify whether there are differences in spectral reflection between conventional and transgenic soybean cultivars RR1 and RR2 under the effect of glyphosate herbicide.

## Material and Methods

### Cultivation of plant material

The experiment was carried out in a greenhouse at the Department of Agriculture of the Federal University of Lavras (UFLA), state of Minas Gerais, Brazil. Six soybean cultivars were used, in which two conventional cultivars (DM-111, DM-Nobre), two cultivars carrying the glyphosate resistance gene (6301, 96Y90) from generation one (RR1) and two cultivars carrying the glyphosate resistance gene (95R40-IPRO, 96R29) from generation two (RR2). The experimental design was a complete randomized block design with five replications. Sowing was carried out in 5-liter pots with soil and sand substrate in a 2:1 ratio, in which five plants per pot were kept. The sowing was carried out on October 21, 2022. Fertilization was carried out according to the recommendation for pots by (MALAVOLTA, 1980) and (NOVAIS; NEVES; BARROS, 1991). At the sowing, an inoculation with *Bradyrhizobium japonicum* and seed treatment with TopSeed® and Cropstar® were performed, as well as other management practices carried out according to monitoring and crop requirements.

### Evaluation of the leaf senescence in trays

To evaluate the effect of the herbicide glyphosate on the plants, a trial was carried out with five treatments, with doses of 0.00, 0.03, 0.06, 0.12 and 0.24% of acid equivalent (AE) of the commercial product Roundup Transorb R. The trial was carried out at the Laboratory of Plant Phenomics at the Department of Biology in the UFLA. Three random leaves of each cultivar were detached 32 days after sowing, in which each leaf represented a repetition. Thus, the experimental design was completely randomized, with three replications in a 6 × 6 factorial scheme (doses per cultivar). The detached leaves were subjected to a herbicide solution soaked in cotton at the base of the detached leaf, in plastic trays with a bottom coated with Germitest paper, so that the solution at different concentrations corresponded to each treatment, generating constant exposure to the product. The leaves were kept under artificial light with 16 hours of light and 8 hours of darkness. Leaf senescence was monitored daily.

### Obtaining the SPAD index

Indirect measurements of leaf chlorophyll were performed with the SPAD-502 (Konica-Minolta, Japan, display range from -9.9 to 199.9 SPAD units) in three areas, two on the sides and the edge of the leaf blade, freeing the midrib. The collection was carried out 14 days after incubation (DAI) in the herbicide.

### Collection of RGB images and Information extraction of from leaves

The collection of RGB images from the leaves was carried out using a digital smartphone camera inside a photo studio to preserve the quality, homogeneity of lighting and pixel size, located at the Laboratory of Plant Phenomics of the Department of Biology at UFLA. In possession of the RGB images, these were then submitted to the section of the useful area that contains the leaves in the image and then submitted to several Python scripting for segmentation using manual thresholding and the RGB, YCrCb, Lab and HLS color spaces to remove the background of the image. Next, after carring out the segmentation and croping of each isolated leaf, the extraction of the average in the red digital number (RDN), green digital number (GDN) and blue digital number (BDN). This procedure was performed using the OpenCV library of the Python language (PYTHON SOFTWARE FOUNDATION, 2022).

### Excess red color index calculation

In order to increase the discrimination of the information in the RDN of the image, the *Excess Red* (ExR) index was adapted from Meyer, Hindman and Laksmi (1999), as in the equation one.

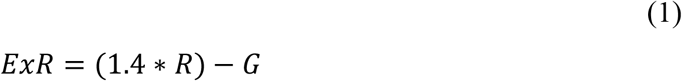

In which:

*ExR*: Corresponding to *Excess Red* index;

*R*: RDN;

*G*: GDN.

### Statistical data analysis

The data obtained were submitted to a descriptive analysis, and for outliers identification. The response variable were assessed according to the following statistical model:

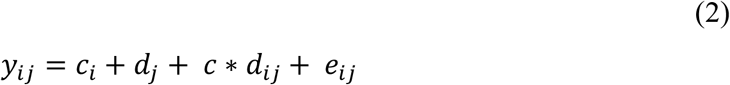

In which:

*y*_*ij*_: value of the observation related to the effedt of the cultivar *i* and the dose *j*;

*c*_*i*_: effect of the cultivar *i*;

*d*_*j*_: effect of the dose *j*;

*c* * *d*_*ij*_: effect of the interaction between the cultivar *i* and the dose *j*;

*e*_*ij*_: error effect associated to the observation of the cultivar *i* and the dose *j*.

Significant effects for cultivar were analyzed using the Skott-Knott test at the 0.05 significance level and dose effects were adjusted for linear regression models. Mean treatment values for each variable were submitted to Spearman’s correlation analysis to assess the relationship between them, and the values were analyzed using the t-test at the 0.05 significance level. Data were analyzed using the R software (R CORE TEAM, 2020).

## Results

### Glyphosate concentrations affect the color traits of soybean cultivars

The chlorophyl activity of the plants is affected by several intracellular factors. The partiotioning of the RGB digital images and obtening the ExR index and the SPAD index were successful in determining of the activity of chlorophyl under the effect of glyphosate in the soybean cultivars studied. The summarized results of the analysis of variance are shown in Table 1 for the variables SPAD and ExR indexes and the R, G and B digital numbers (respectively RDN, GDN, and BDN). There was a significant effect only for the glyphosate herbicide factor. The soybean cultivars showed no difference in light reflectance under the effect of the herbicide and there was also no significant effect of the interaction between the factors.

**Table 1.** Mean square (MS) of the SPAD and ExR indexes and average digital numbers for the channels R, G and B of the soybean cultivars submitted to different doses of glyphosate applied to the dettached leaves under lab conditions. Federal University of Lavras, Lavras, state of Minas Gerais, Brazil, 2022.

| MS | SPAD | ExR | RDN | GDN | BDN |
| --- | --- | --- | --- | --- | --- |
| Cultivar | 86.68 | 1765.60 | 780.40 | 745.10 | 96.60 |
| Doses | 717.38* | 11353.25* | 6456.50* | 4194.90* | 3756.90* |
| C*D | 61.44 | 983.50 | 493.50 | 3003.80 | 151.40 |
| Error | 47.46 | 1136.23 | 553.67 | 8568.80 | 1405.10 |
| CV (%) | 19.70 | 54.71 | 32.07 | 10.58 | 11.77 |

\* Significant at the level of 0.05 by the *F* test.

The data related to the SPAD and ExR indexes and the digital numbers of the three color channels of the RGB space are described in Table 2. There was extensive variation for the SPAD index, raging from 8.6 to 59.4, and the higher the value indicates that higher is the chlorophyll content in the leaf. The ExR index ranged from 14.25 to 170.92. Regarding the digital numbers, the RDN ranged from 33.44 to 155.35, the GDN ranged from 84.16 to 139.12, and for BDN, from 22.92 to 57.18 as a function of senescence variation generated by the effect of herbicide doses. The ExR index is used to increase the differentiation of the effect of RDN in regard to the others to minimize interference from the other channels in the image.

**Table 2.** Distribution of the values of the variables SPAD and ExR indexes and the average R, G and B digital numbers of the soybean cultivars submitted to different doses of glyphosate applied to leaves dettached under lab conditions. Federal University of Lavras, Lavras, state of Minas Gerais, Brazil, 2022.

| Mean | SPAD <sup>a)</sup> | ExR | RDN | GDN | BDN | Virtual Color <sup>b)</sup> |
| --- | --- | --- | --- | --- | --- | --- |
| Minimum | 8.60 | 14.25 | 33.44 | 84.16 | 22.92 |  |
| 1<sup>st</sup> quartile | 30.80 | 31.62 | 55.09 | 102.07 | 36.21 |  |
| Median | 38.15 | 47.46 | 65.49 | 111.53 | 41.10 |  |
| Average | 34.97 | 61.61 | 73.37 | 112.99 | 41.11 |  |
| 3<sup>st</sup> quartile | 40.20 | 90.68 | 89.57 | 125.04 | 45.01 |  |
| Maximum | 59.40 | 170.92 | 155.35 | 139.12 | 57.18 |  |

<sup>a)</sup> SPAD index ranges from -9.99 to 199.99. The higher the SPAD values, the higher the chlorophyll content.
<sup>b)</sup> Virtual color obtained by combining the values of each channel after splitting R, G, and B channels. Values range from 0 to 255 in the RGB color space.

The doses of glyphosate herbicide caused contrasting effects in all studied variables. In the literature it is possible to identify that the SPAD index correlates directly with the chlorophyll content (r = 0.88) in the soybean leaf (GWATA, 2004). In this study, the doses caused a progressive decrease in the SPAD index of soybean leaves, as shown in Figure 1.

**Figure 1.**
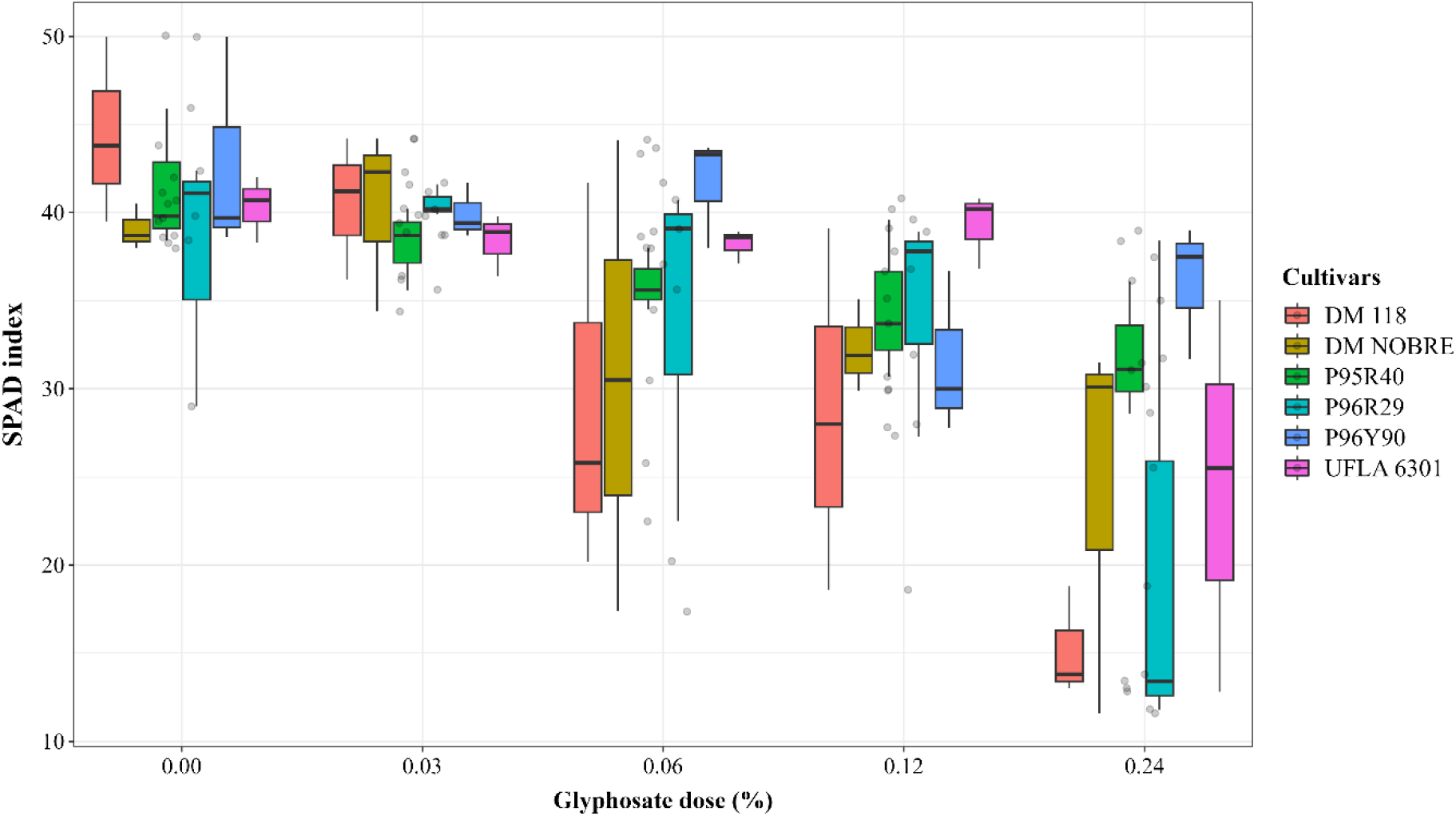
Boxplot diagrams of the SPAD index for the glyphosate doses, discriminating the effect of the cultivars. Federal University of Lavras, Lavras, state of Minas Gerais, Brazil, 2022.

The minimum SPAD value presented by the leaves at the maximum dose was 8.6. In the leaves of the control treatment, the average SPAD value was 41.43, corresponding to a 4.8-fold reduction. In general, the increasing doses of glyphosate reduced the chlorophyll content, indicating the occurrence of intoxication by the herbicide even in RR cultivars, due to the increase in dose and continuous exposure. In the other hand, the increasing doses of glyphosate promoted an increase in the reflectance of the RDN, directly reflecting a reduction of absorption of this color band (Figure 2). The average of the control treatment was 75.85 and the treatment 0.24% AE dose was 66.64. The average value of the RDN presented by the dose recommended for the soybean crop (0.06% AE) was 74.27. The 0.03% dose showed a drop in relation to the control treatment. The R channel is the main absorbed sunlight, being the most used by the antenna complex, which is composed of photosystems I and II (TAIZ et al., 2017), and greater the reflection of the R channel, greater the damage for photosynthesis.

**Figure 2.**
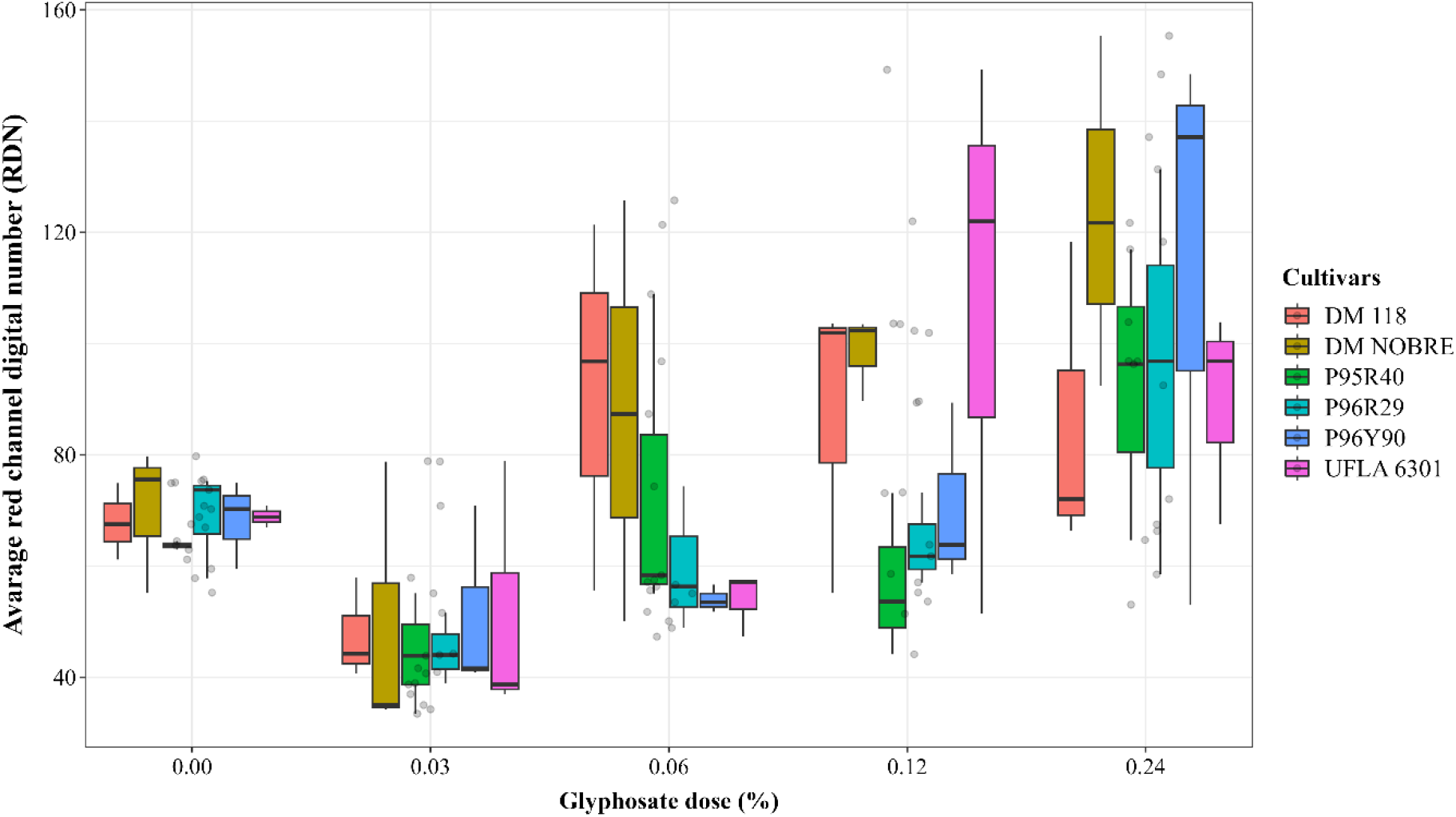
Boxplot diagrams for the average RDN from the RGB color space according to the glyphosate doses, differentiating the effects of the soybean cultivars. Federal University of Lavras, Lavras, state of Minas Gerais, Brazil, 2022.

Figure 3 shows the effect of glyphosate herbicide doses for each cultivars. A similar behavior is verified with increasing doses for all cultivars, with the linear model being the one that best adjusted the data variation.

**Figure 3.**
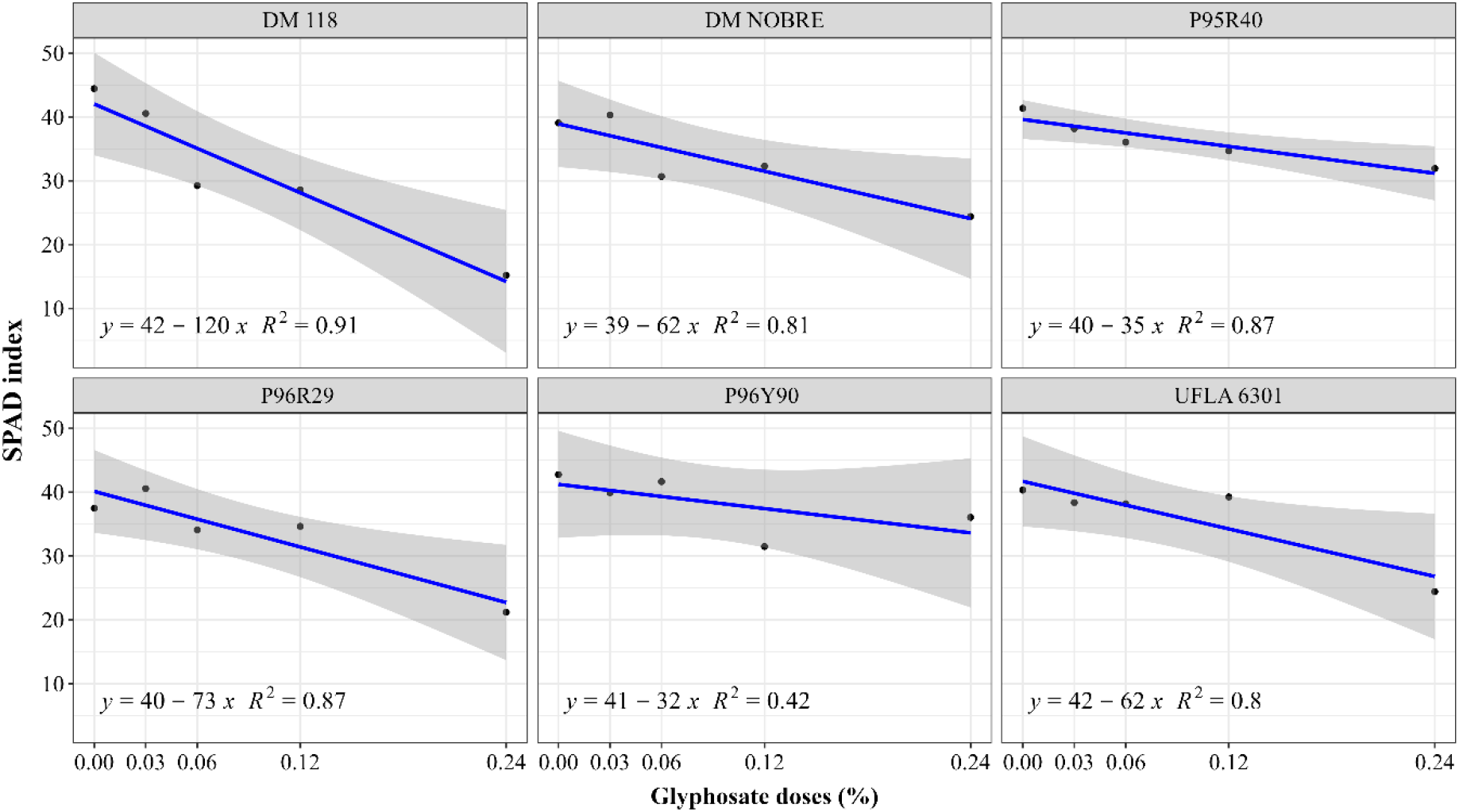
Linear regression analysis for each cultivar studied for the SPAD variable of the RGB color space as a function of glyphosate doses. Federal University of Lavras, Lavras, state of Minas Gerais, Brazil, 2022.

The greatest angle of the straight line, represented by the coefficient β1 of the regression equation, was obtained for the conventional cultivar DM 118 (120-fold decrease) and which obtained the highest coefficient of determination (R^2^ = 0.91); however, there is a high inclination towards to cultivar P96R29 (Figure 4). This result indicates that this cultivar has a lower tolerance to high doses of the herbicide compared to the others, which may result in greater damage to the treated plants. Regarding the results of regressions for each cultivar for the average RDN as a function of glyphosate doses, as previously shown, there was an inverse relationship with the SPAD index for all cultivars; however, the highest reflectance for this band was obtained by the conventional cultivar DM Nobre, while the lowest for DM 118. This result refleects the fact that these values refer to the leaf area as a whole, regardless of the necrosis generated by the action of the herbicide, so it is appropriate to isolate the area to study the specific reflectance of the specific leaf area.

**Figure 4.**
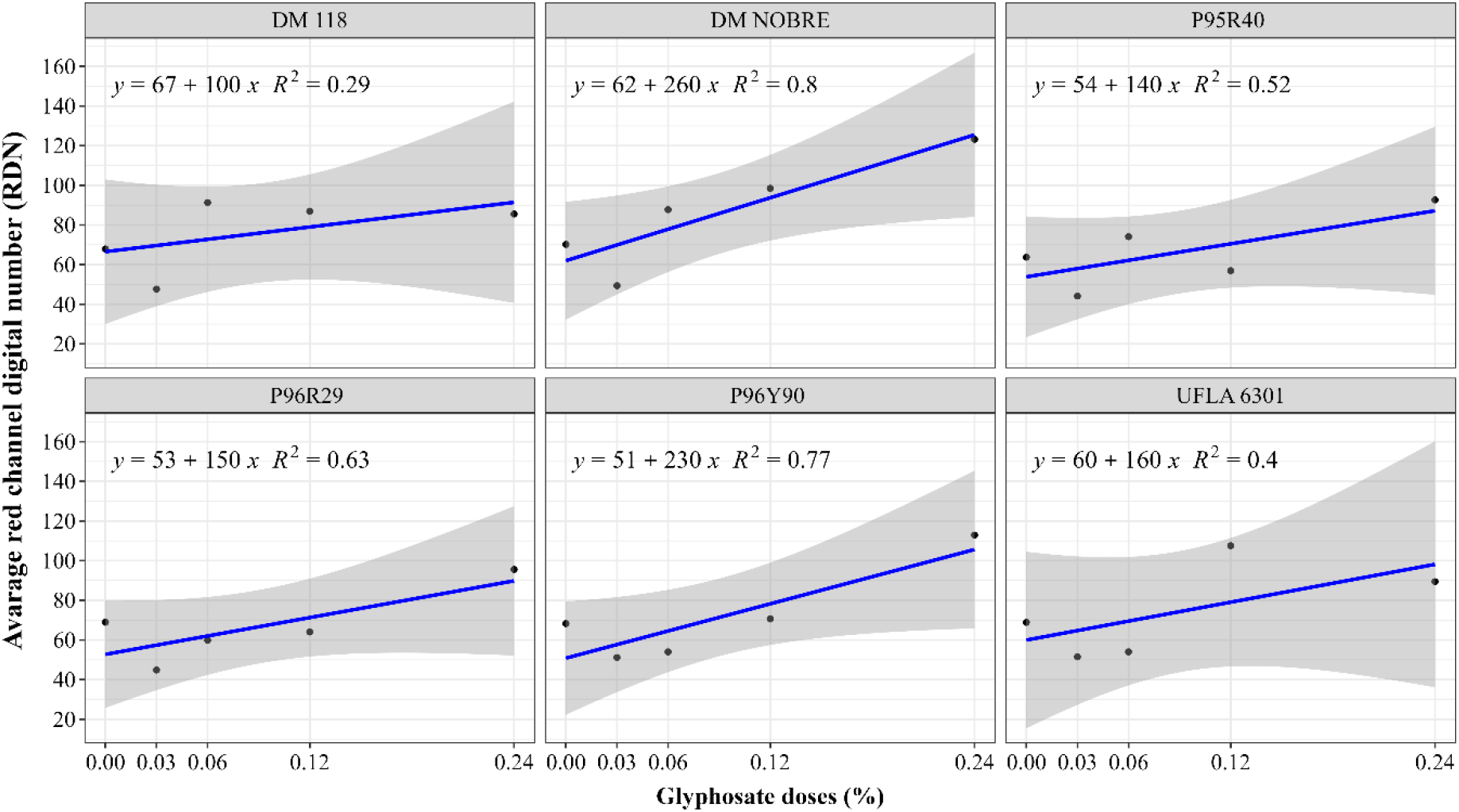
Linear regression analysis for each cultivar studied for the average RDN of the RGB color space as a function of glyphosate doses. Federal University of Lavras, Lavras, state of Minas Gerais, Brazil, 2022.

Regarding the average GDN and BDN, respectively shown in Figures 5 and 6, the responses due to the increase in the herbicide dose were low for all cultivars without a defined pattern, which highlights the importance of the R channel as a potential variable related to the response to the effects of glyphosate in conventional and resistant soybean cultivars.

**Figure 5.**
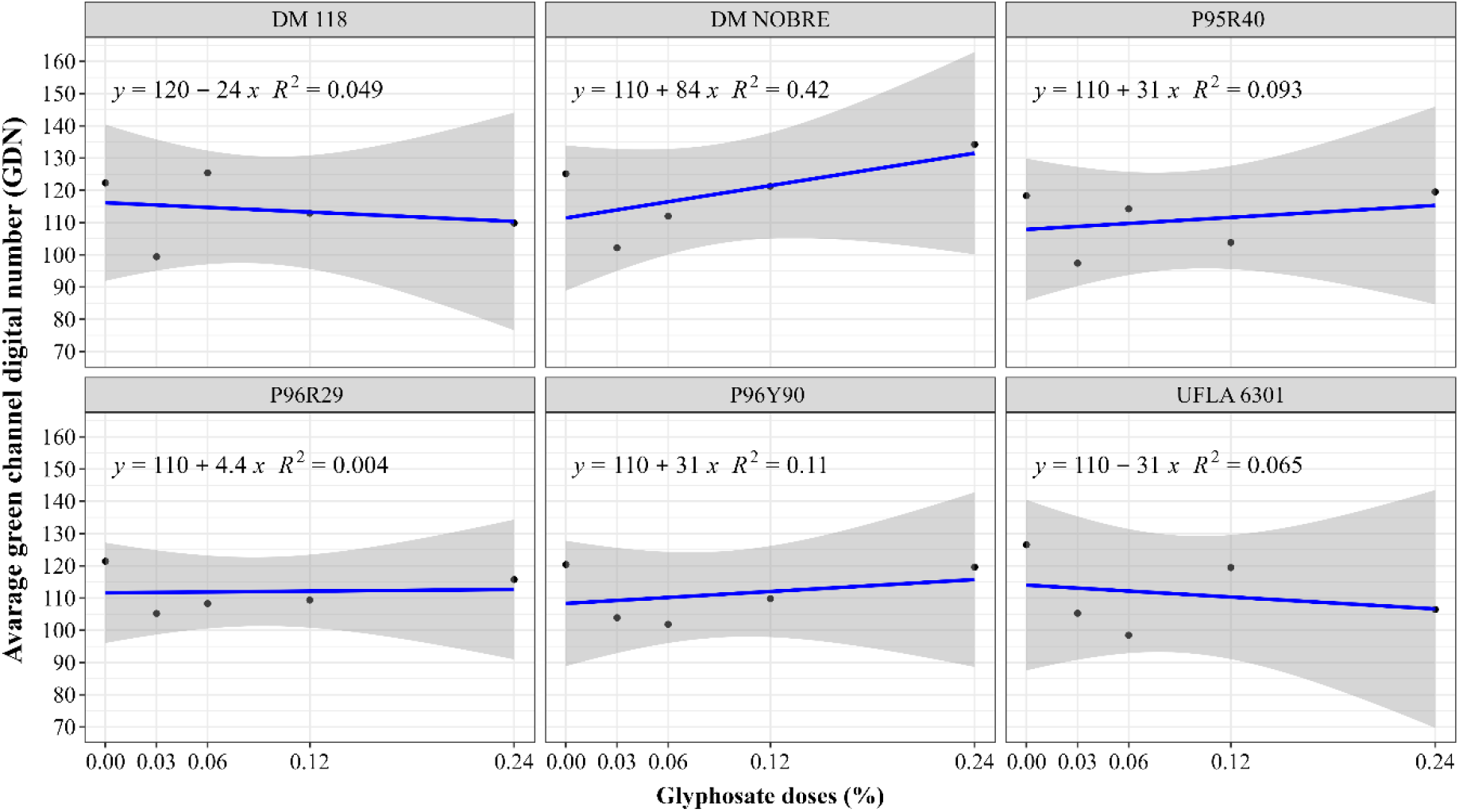
Linear regression analysis for each cultivar studied for the average GDN of the RGB color space as a function of glyphosate doses. Federal University of Lavras, Lavras, state of Minas Gerais, Brazil, 2022.

**Figure 6.**
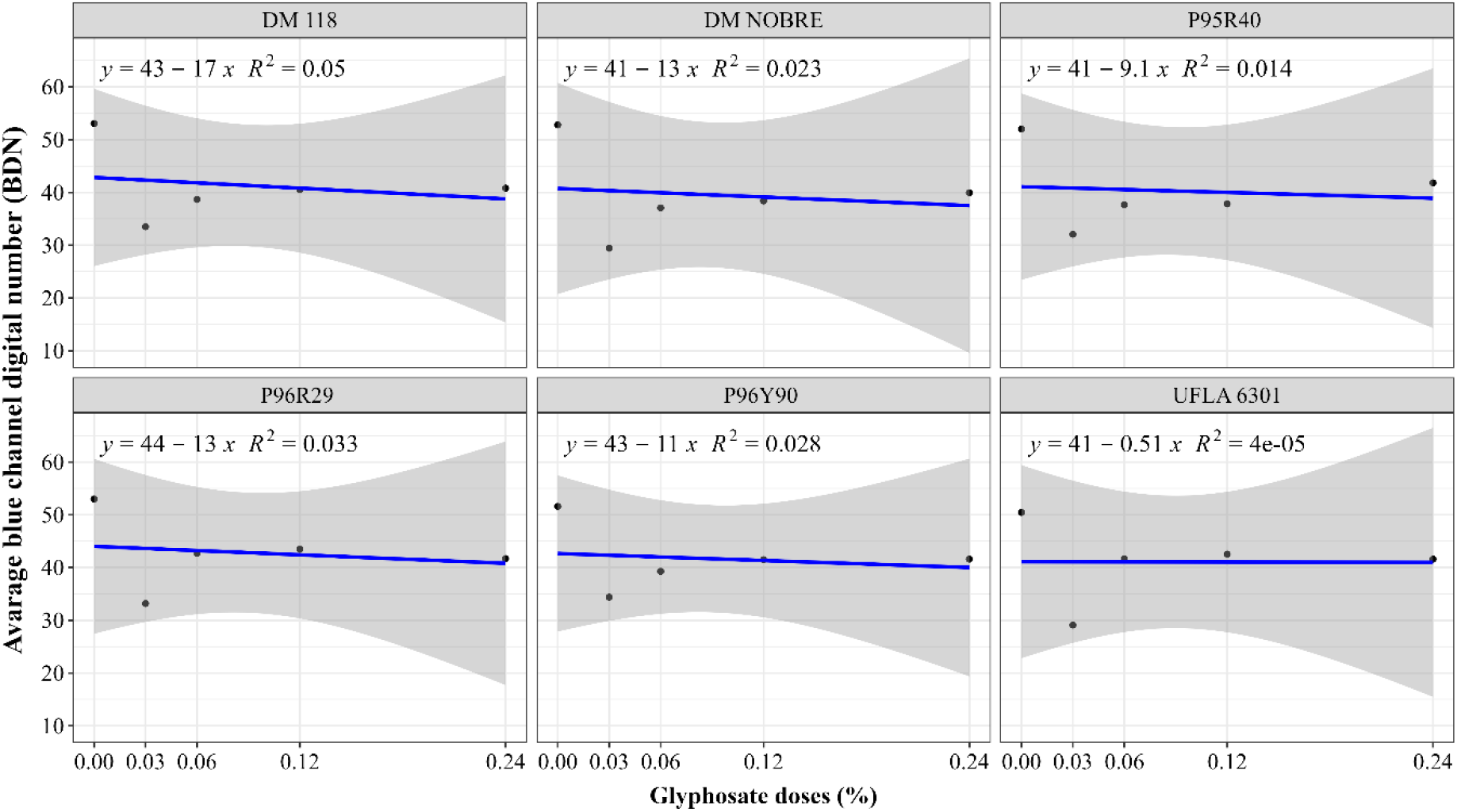
Linear regression analysis for each cultivar studied for the average variable of the average BDN of the RGB color space as a function of glyphosate doses. Federal University of Lavras, Lavras, state of Minas Gerais, Brazil, 2022.

Due to this low variation observed in the average GDN and BDN, it was possible to isolate the effect of red channel effect directly, being highlighted by the application of the ExR index (Figure 7), which resulted in behavior similar to that observed for the RDN for each cultivar studied.

**Figure 7.**
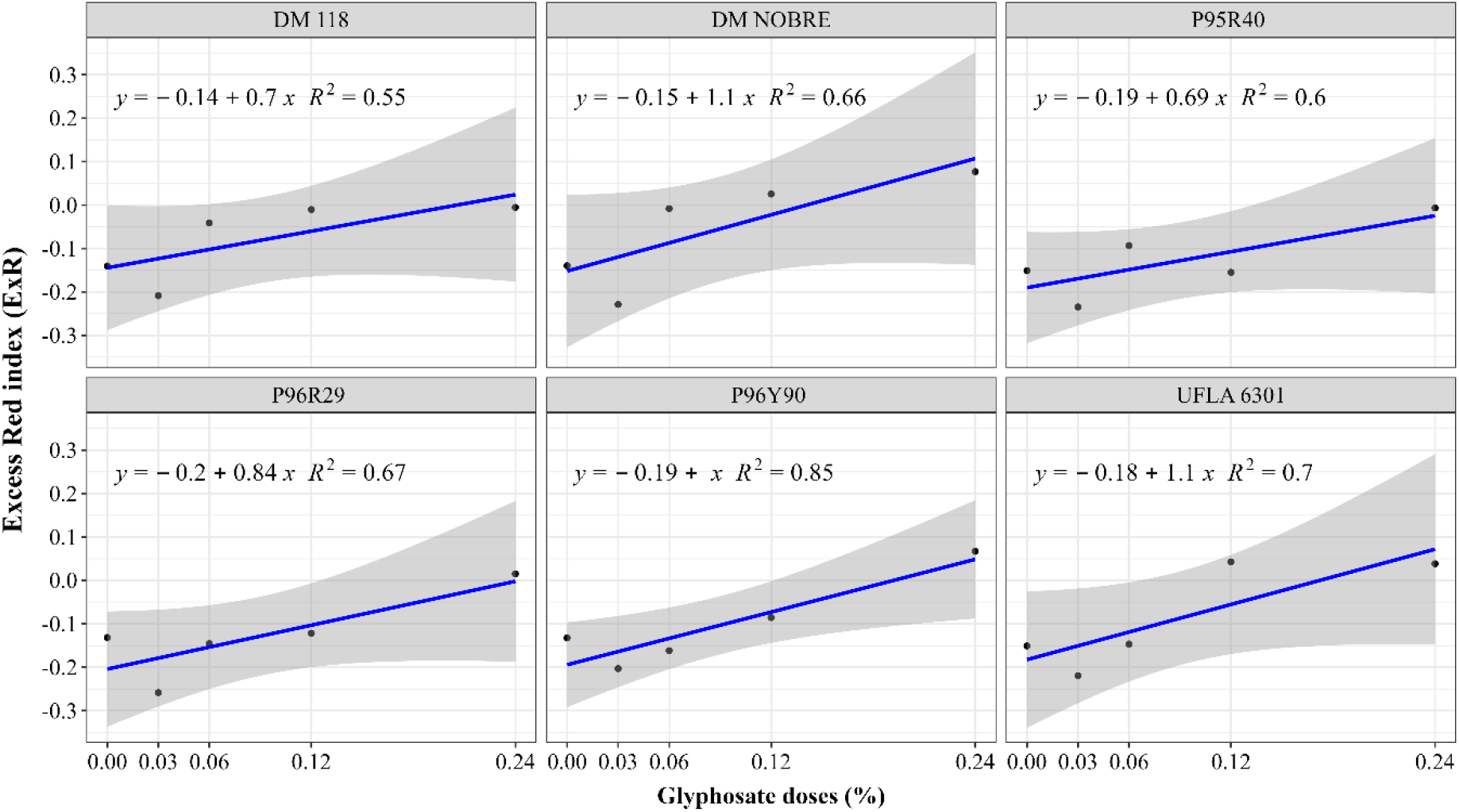
Linear regression analysis for each cultivar studied for the ExR index as a function of glyphosate doses. Federal University of Lavras, Lavras, state of Minas Gerais, Brazil, 2022.

An important characteristic obtained with the use of the ExR index was a better precision of the coefficient of determination, which makes the inferences more accurate about the cultivars in relation to the isolated information of the average RDN, in which the values of R^2^ ranged from 0.29 to 0.80, and for the ExR from 0.37 to 0.89. These variables present a significant, direct and high relationship of 0.97 according to Spearman’s correlation analysis (Figure 8). This indicates that the average RDN alone is sufficient to represent the herbicide effect on the red color reflectance. There was also a significant and high correlation (>0.58) between the GDN and RDN, GDN and ExR, and BDN and GDN. The SPAD index also showed a high but negative correlation with the RDN (−0.62) and with the ExR index (−0.66). The mentioned significant correlations prove that the herbicide effect really interfered in the reflectance of the RDN and reduced the chlorophyll content in the leaf through the SPAD index.

**Figure 8.**
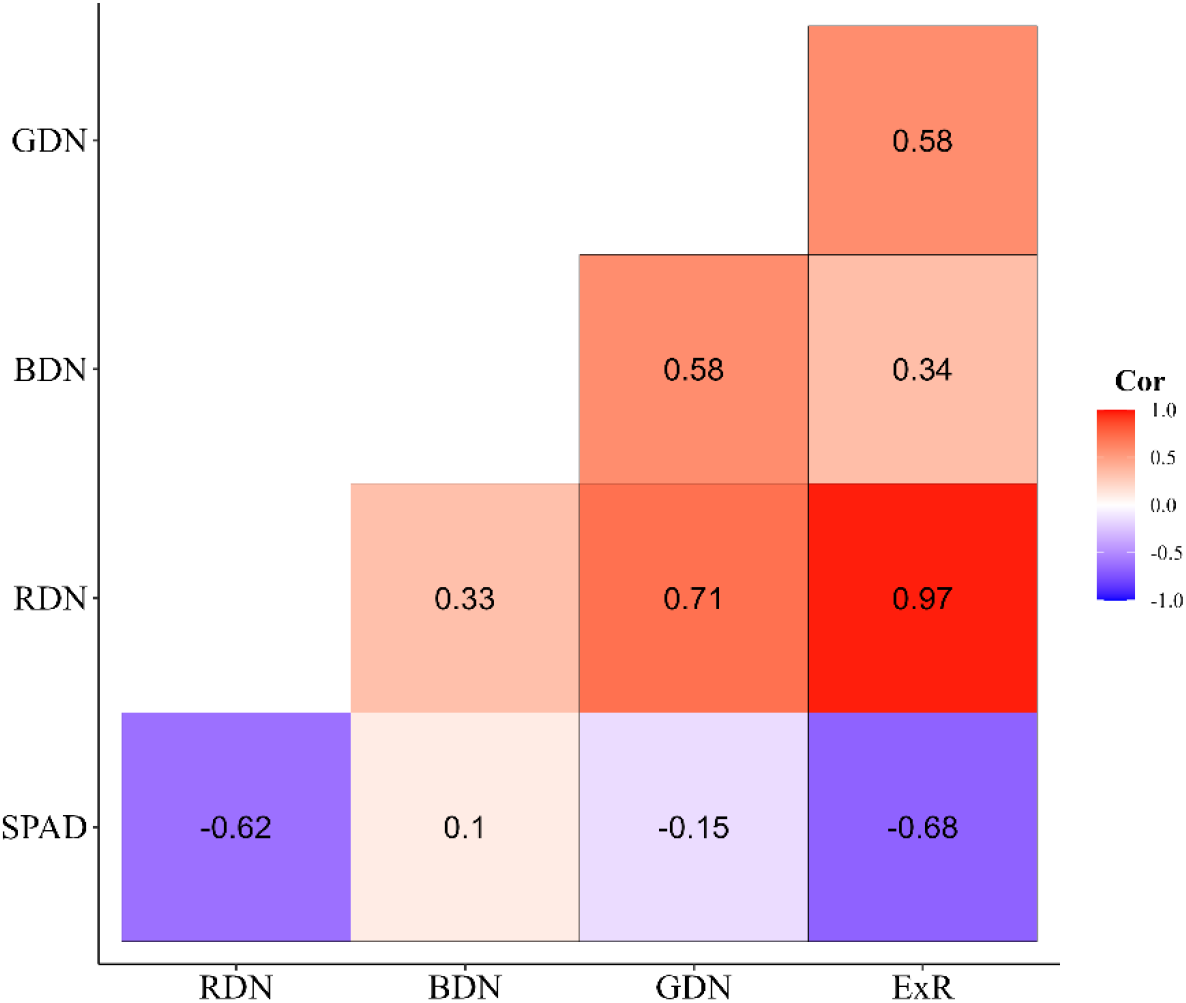
Spearman’s correlation coefficients of the variables studied in soybean cultivars subjected to different doses of glyphosate applied to detached leaves under controlled conditions. Federal University of Lavras, Lavras, state of Minas Gerais, Brazil, 2022. * The values presented are significant at 0.05 level by the t-test. Cor: Spearman’s correlation.

## Discussion

### Leaf phenotyping with smartphone camera

Digital images obtained with a smartphone camera are a great tool for phenotyping soybean leaves. Previous studies have already shown that this type of digital camera can be used to perform the task of identifying colors with quality in the field (HASSANIJALILIAN et al., 2020) and under controlled conditions, such as a greenhouse (RIGON et al., 2016), showing how an added advantage is obtaining a large amount of data from few images. Our study was carried out in a similar way, with the plants being conducted in a greenhouse and the images obtained in the studio in order to control the variation in lighting. Our study is the first contribution that shows the effect of the glyphosate herbicide on the reflectance of visible light, and correlating the values of the SPAD indix and the digital channels of the RGB space. Our study paves the way for further research related to the effects of agricultural chemicals on the photosynthetic apparatus of plants by using digital images.

### Effect of absorption of glyphosate herbicide

The glyphosate herbicide acts by blocking the 5-enolpyruvylshikimate-3-phosphate synthase (EPSPS) route in both mono and dicots (STEINRÜCKEN; AMRHEIN, 1980), an enzyme that performs the metabolism of 3-phosphoshikimate into 5-enolpyruvylshikimate-3-phosphate, which is a fundamental step for the synthesis of aromatic amino acids, such as tryptophan, phenylalanine and tyrosine, which leads to a total shortage of these essential amino acids and plant death (SCHÖNBRUNN et al., 2001). Surprisingly, our results proved that this herbicide affects the absorption of red light, increasing its reflectance and causing damage to photosynthesis, preventing its absorption by the antenna complex. This is a problem to be analyzed, because with frequent applications of this herbicide per cycle of the soybean crop, it will certainly have effects on the expected productivity, even in soybean cultivars that contain the resistance gene.

In addition, the application of excess volume of glyphosate has increased the cases of weeds with tolerance to the herbicide. The unconscious use of doses such as glyphosate, as well as non-compliance with technical recommendations, has increased the number of plants resistant to this herbicide. In plants of the *Amaranthus tuberculatus* species, eight additional copies of the EPSPS gene have been reported, resulting in amplification of the expression of this regulatory gene, resulting in tolerance to the acidic ingredient. Yet in this study, no point mutation was observed for amino acids G101, T102 or P106 in the coding sequence by the EPSPS gene (LORENTZ et al., 2014), which endorses the tolerance of these plants to the acidic ingredient.

## Conclusion

Glyphosate affects the absorption of red light in resistant plants when subjected to continuous exposure, leading to senescence and that the red channel information can be used to infer the level of interference in the photosynthetic activity of plants subjected to the herbicide.

